# PADI4-mediated citrullination of histone H3 stimulates HIV-1 transcription

**DOI:** 10.1101/2024.03.17.583304

**Authors:** Luca Love, Bianca B Jütte, Birgitta Lindqvist, Hannah Rohdjess, Oscar Kieri, Piotr Nowak, J Peter Svensson

## Abstract

HIV-1 infection establishes a reservoir of long-lived cells with integrated proviral DNA that can persist despite antiretroviral therapy (ART). Some of these reservoir cells can at anytime be reactivated and reinitiate infection. The mechanisms governing proviral latency the transcriptional regulation of the provirus are complex and have not yet been sufficiently described. Here, we identified a role for histone H3 citrullination, a post-translational modification catalyzed by protein-arginine deiminase type-4 (PADI4), in HIV-1 transcription and latency. We found that PADI4 inhibition by GSK484 reduced HIV-1 transcription after T cell activation in *ex vivo* cultures of CD4 T cells from people living with HIV-1 (PLWH). The effect was more pronounced in individuals with active viremia compared to individuals with effective ART. Using cell models of HIV-1 latency, we showed that PADI4-mediated citrullination of histone H3 occurred at the HIV-1 promoter upon T cell stimulation which facilitated proviral transcription. HIV-1 preferentially integrated into genomic regions marked by H3 citrullination and these integrated proviruses were less prone to latency compared to those in non-citrullinated chromatin. Inhibiting PADI4 led to compaction of the HIV-1 promoter chromatin and an increase of HP1α-covered heterochromatin, in a mechanism partly dependent on the HUSH complex. Our data reveal a novel mechanism to explain HIV-1 latency and transcriptional regulation.

**Highlights:**

- The PADI4 enzyme stimulates HIV-1 transcription during T cell activation.
- PADI4 citrullinates histone H3 at the HIV-1 promoter upon T cell activation and inhibiting PADI4 reduces HIV-1 reactivation in *ex vivo* CD4 T cells from people living with HIV-1.
- H3cit is mostly found at gene promoters, and productive HIV-1 proviruses are more likely than latent or reactivatable proviruses, to integrate in chromatin susceptible for citrullination.
- H3cit inhibits latency establishment by interfering with the binding of HP1α to H3K9me3, preventing heterochromatin formation.

## Introduction

HIV-1 establishes a latent viral reservoir that persists despite active antiretroviral therapy (ART). The cells in the latent reservoir represent a heterogeneous population in different T cell subsets. Even though the cells in the latent reservoir do not produce viral particles, the HIV-1 provirus can still be transcriptionally active ^1–3^. In other cells, the provirus is likely to be permanently silent, such as those integrated in epigenetically silent regions ^4–6^. Before the start of ART, HIV-1 gradually depletes the CD4 T cell population. However, once ART is initiated, latently infected cells are selected for, since the host immune system eliminates cells expressing viral epitopes ^7^. Some of these latent cells can switch between silent and active states of the provirus to avoid detection by the immune system, thus maintaining the ability to re-initiate active infection if ART is interrupted. Latency can be induced by many factors (reviewed in ^8^), but a key determinant in long term latency is the chromatin constitution of the provirus ^5,6^. As HIV-1 enters the genome, it is assumed to take the epigenetic profile of the chromatin where it integrates. However, the host cell can stably silence a locus through different mechanisms including the KRAB-ZFP (KAP1) transcription factor and Human Silencing Hub (HUSH) ^9^ leading to heterochromatin protein 1 (HP1) deposition over the provirus ^10,11^. A more transient proviral inactivation is provided by lysine deacetylations ^12^.

The proviral sequence resembles a host gene in several aspects, and the resulting transcripts are initially processed similarly to human mRNAs. However, at the stage of virus production, the spliceosome is turned off and the unspliced viral RNA genome is transcribed to be packaged into virions^13^. It is unclear what promotes this switch, but acetylation of viral transactivator protein Tat has been proposed as one mechanism leading to loss of splicing ^14^. Unspliced transcripts are recognized as foreign by the HUSH complex, and specifically the TASOR subunit. This recognition initiates a process of silencing the chromatin whereby the histone H3 lys9 is methylated (H3K9me3) to enable heterochromatin ^15^.

A chromatin modification that has been associated with preventing silencing of foreign DNA ^16^, yet not studied from the perspective of HIV-1, is histone citrullination. Histone citrullination affects the compaction of nucleosomes ^17^, thereby affecting chromatin accessibility of promoters ^18^. These changes in chromatin accessibility have been attributed to the reduced electrostatic attraction between histones and DNA when positively charged arginine is converted to neutral citrulline ^17^. Histone modifications serve as a platform for the binding of other proteins. H3K9me3 is recognized and bound by the chromodomain of the α variant of HP1 (HP1α) ^19^. Modification of arginine adjacent to lysine residues can affect the binding of chromodomain proteins that recognize methylated lysines^16^. Arginines can be modified by methylation or citrullination ^20^. Citrullination is mediated by members of the protein-arginine deiminase (PADI) family, of which PADI2 and PADI4 are found in the nucleus and have partially redundant functionality ^21^. PADI4 is an enzyme primarily expressed in less differentiated cells and various cancer cells ^22–25^. It citrullinates targets in the cell nucleus, including several residues of histone H1, histone H3 and HP1γ ^25–27^. H3cit8 interferes with HP1α-binding to H3K9me3 in the process of regulating human endogenous retroviruses (HERVs) ^16^.

Here we describe a PADI4-mediated citrullination effect on HIV-1 transcription and latency reversal. *De novo* histone H3 citrullination accompanies HIV-1 transcription during cell activation and prevents proviral latency establishment, especially before ART is initiated. PADI4 acts partly through H3R8cit that interferes with HIV-1 latency initiation by interfering with the binding of HP1α to H3K9me3, preventing heterochromatin formation at the HIV-1 promoter.

## Results

### PADI4 stimulates HIV-1 transcription after cell stimulation in cells from viremic ART naïve people with HIV-1

To test the hypothesis that citrullination affects the latent HIV-1 provirus in reservoir cells from PLWH, we recruited 31 study participants to a clinical study. Given the known effect of PADI4-mediated histone citrullination on heterochromatin ^16,25^ together with the observation that the reservoir of intact proviruses becomes heterochromatic with time ^6,28^, we included two groups in our cohort. Firstly, viremic study participants (n=14) and secondly, participants with long-term (>6 years) ART suppressed viremia (<50c/ml) (n=17). The study participants were matched across various characteristics, apart from their plasma viremia levels and ART status (Table S1). CD4 T cells were isolated from peripheral blood. The cells were activated by phorbol-12-myristate-13-acetate and ionomycin (PMAi) after treatment with PADI4 inhibitor GSK484 or mock treatment. The intact HIV-1 reservoir was measured using Intact proviral DNA assay (IPDA) ^29^. Cell-free RNA was measured by the amount of intact virus RNA in the supernatant ^30^. For cell-associated RNA, the LTR region and the multiply spliced RNA was investigated.

In the entire cohort, GSK484 reduced the *ex vivo* HIV-1 activation based on cell-free RNA (Fig 1A). RNA data could be obtained from 21 study participants (viremic: n=9, ART: n=12). Stratification between viremic and ART suppressed study participants showed that cell activation induced a release of intact viral particles only in cells from viremic study participants. The virus release was dependant on PADI4 as it was abolished in cells treated with GSK484. Activation of cells from the ART treated study participants did not result in release of intact viruses (Fig 1B). This difference was not a consequence of different numbers of infected cell as the intact HIV-1 reservoir was similar between the two groups (Fig 1C). We continued to investigate the RNA prior to viral release. The cell-associated RNA from the LTR transcripts showed that both groups had a clear proviral activation after PMAi-stimulation (Fig 1D). This increase of LTR transcripts was not significant in the presence of GSK484. A comparison between PMAi-stimulated cells showed a significant reduction in viremic samples pre-exposed to GSK484 (Fig 1D). Multiply spliced processed RNA (Tat-Rev) was induced by PMAi activation in cells from ART suppressed study participants only (Fig 1E), implying involvement of Tat in the latency reversal process in this group. A lack of mRNAs encoding Tat in activated cells from viremic study participants suggests that latency reversal in these cells was Tat-independent or that the Tat protein was already present in these cells. As the effect of GSK484 was most strongly associated with viremic study participants, we quantitated the levels of *PADI4* to determine if the differences could be attributed to different protein amounts. We observed that the expression of *PADI4* varied considerably in the cohort, but there was no difference between *PADI4* expression in resting CD4 T cells from viremic or ART suppressed study participants (Fig 1F).

**Fig 1:**
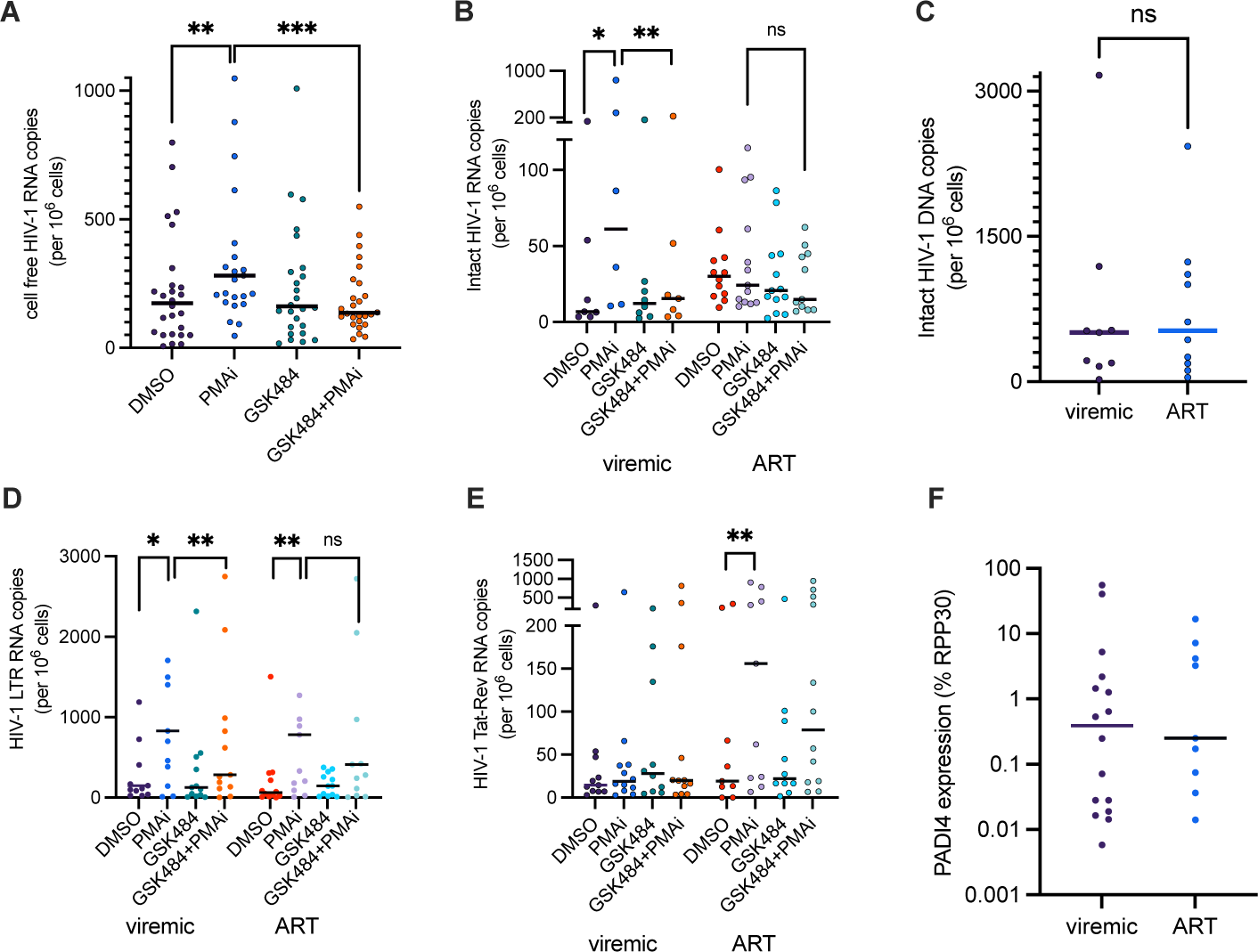
PADI4 stimulates HIV-1 transcription after cell stimulation in cells from viremic people with HIV-1. (A) Extracellular HIV-1 RNA with PMAi and GSK484 treatments from entire cohort (n=31, study participant characteristics in Table S1). (B) Intracellular cell-associated HIV-1 RNA in the cohort stratified into in viremic and long-term ART suppressed groups. (C) Intact HIV-1 proviral reservoir from the two groups measured by IPDA. (D-E) HIV-1 5’LTR RNA (D) and Tat-Rev (E) quantified in viremic (n=14) and long-term ART suppressed groups (n=17). (F) PADI4 expression in the two groups normalised to RPP30. All statistical tests shown are Wilcoxon matched-pairs signed rank test * p<0.05, ** p<0.01, *** p<0.001.

This data show that PADI4 influences HIV-1 latency reversal in cells from people living with HIV-1, particularly before the initiation of ART.

### PADI4 facilitates HIV-1 transcription after cell stimulation in cell models

To investigate mechanisms underlying the effects of PADI4 inhibition on HIV-1 proviral activation, we used established cell models of HIV-1 latency. The J-lat clone 5A8 is a Jurkat-derived cell line with a single copy of a reporter HIV-1 integrated in an intron of the *MAT2A* locus. Proviral expression can be tracked by flow cytometry or microscopy as the sequence for *nef* is replaced by *gfp* ^31,32^. To test the selectivity of PADI enzymes for the effect on HIV-1 transcription, cells were exposed to three different PADI inhibitors: pan-PADI inhibitor Cl-amidine, PADI2-specific CAY10723, and PADI4-specific GSK484. The effect was measured by flow cytometry after 24 h exposure. The PADI inhibitors alone had no effect on HIV-1 latency reversal (Fig 2A). When cells were exposed to PADI inhibitors in combination with PMAi to stimulate the cells, Cl-amidine and especially GSK484 resulted fewer GFP-positive cells (Fig 2A). We also compared the effect of GSK484 with GSK106, an inactive control of GSK484, showing that GSK484 reduces the level of HIV-1 reactivation compared to GSK106 (Fig S1). This suggests that citrullination by PADI4 is part of a HIV-1 latency reversal mechanism. CAY10723 resulted in no significant decrease in HIV reactivation when the cells were activated with PMAi. PADI4 is known to be expressed in neutrophils ^33^, but the gene is also weakly expressed in many other cell types. In our cells, we verified that PADI4 was expressed, although at low basal level (Fig 2B) (1,700 copies/10 ng RNA). Upon cell stimulation, PADI4 expression was increased (Fig 2B). To understand the kinetics of the GSK484 effect, we set up a time course experiment and observed that the PADI4 inhibition led to a lower plateau of proviral activation although the kinetics of latency reversal were similar (Fig 2C). This is consistent with a subpopulation of cells being affected by PADI4 inhibition. To test if the short GSK484 half-life postpones activation, we also performed a pulse experiment with PMAi and GSK484 present for one hour and then washed out. The results were similar between a pulse or continuous exposure indicating that the effect we observed is induced in the first hour of activation, at the early stages of HIV latency reversal (Fig 2C).

**Fig 2.**
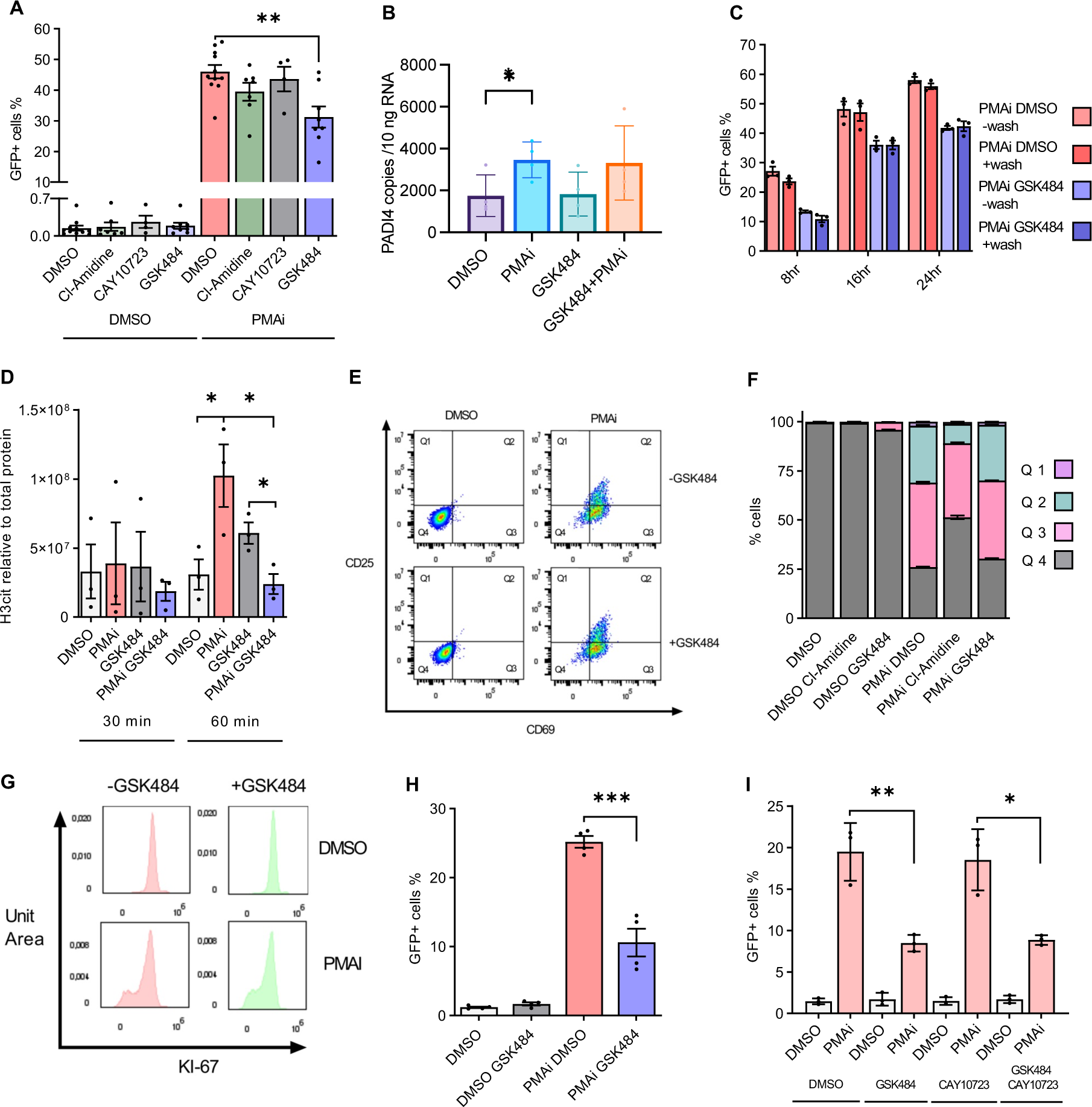
PADI4 facilitates HIV-1 transcription after cell stimulation in cell models. (A) J-Lat 5A8 cells treated with PADI inhibitors and PMAi activation. GFP measured by flow cytometry (n=8-11). (B) PADI4 expression in 5A8 measured by ddPCR under PMAi and GSK484 treatment (n=4). (C) Time course of 5A8 under PMAi and GSK484 treatment and flow cytometry quantification of GFP. Cells were washed and resuspended in new media without drugs after 1 h (n=3). (D) H3cit (R2, R8, R17 residues) quantification by immunoblot in 5A8 with PMAi and GSK484 treatment at two early timepoints (n=3). (E) T cell activation phenotype for 5A8 as measured by flow cytometry. CD69 appears early during T cell activation, CD25 appears late during late T cell activation. (F) T cell activation of 5A8 after 24 h with different PADI4 inhibitors and PMAi activated cells (n=3). (G) Staining with proliferation marker KI-67 of 5A8 treated for 24 h with PMAi and GSK484 (H) Polyclonal K562-Lat treated with PMAi and GSK484 for 24 h and quantified with flow cytometry (n=4). (I) Polyclonal K562-Lat treated with PMAi, GSK484, and CAY10723 for 24 h and quantified with flow cytometry (n=3) Data is shown as mean ± SEM. Statistical tests shown are unpaired T tests * p<0.05, ** p<0.01, *** p<0.001.

In the nucleus, PADI4 citrullinates many targets, including histone H3. As observed previously ^16^, global H3cit was induced within an hour after cell activation as measured by immunoblot. This induction is not observed after PADI4 inhibition (Fig 2D). As expected from the stable nature of the H3cit modification, GSK484 prevents *de novo* citrullination but it does not reduce H3cit in a short time period (Fig 2D).

Stimulating T cells effectively reverses HIV-1 latency. To test if the decreased proviral activation could be explained by GSK484 preventing T cell activation, we determined the abundance of the surface markers CD69 (early activation) and CD25 (late activation). CD25 was not affected by PADI4 inhibition, but for CD69 showed a slight increase after GSK484 exposure (Fig 2E,F). This modest increase in T cell activation is unlikely to explain the reduced HIV-1 transcription, as T cell activation stimulates HIV-1 latency reversal. The GSK484 effect on cell proliferation was measured by Ki-67 expression, which remained unaffected by PADI4 inhibition (Fig 2G).

To confirm a more general role of PADI4 in HIV-1 latency reversal, we inhibited PADI4 by GSK484 in a K562-derived HIV-1 latency model ^34^. In K562-lat cells, the PADI4 gene was expressed (48 copies/10 ng RNA), although at a lower level than in 5A8 of (1800 copies/10 ng) (Fig 2A). This K562-lat cell line is polyclonal with respect to HIV-1 integration sites. As observed for the 5A8 cell line, PADI4 inhibition resulted in reduced PMAi-induced HIV-1 activation (Fig 2H). Inhibiting PADI2 with CAY10723 had no effect alone and no additive effect was observed when inhibiting both PADI2 and PADI4 (Fig 2I), suggesting non-redundancy between the two proteins with respect to HIV-1 latency reversal.

### Clonal effects of T cell activation and PADI4 inhibition

To test the effect of chronic PADI4 depletion, we attempted to mutate PADI4 by CRISPR. We failed to generate homozygous PADI4-/-mutants and heterozygous mutants expressed wild-type levels of PADI4. However, monoclonal selection revealed that cultures recently derived from a single cell had two different phenotypes with regard to PMAi-induced activation and GSK484 treatment (Fig S2A). Whereas half of the clones responded as the bulk culture to T cell activation (30-40%) and GSK484 treatment (23% reduction), half of the clones where easily activated by PMAi (60-80%) and showed a smaller (9%) non-significant effect of GSK484 treatment. Continuous propagation of the cultures reduced this effect. Moreover, the abundance of CD69 marker on the cell surface was significantly increased after GSK484, although not comparable to the CD69 levels after T cell activation (Fig S2B). The clones are expected to be genotypically identical, but as we have showed previously, the provirus in 5A8 bulk culture is epigenetically heterogeneous ^35^. The bimodal phenotype could possibly be explained by the chromatin contexts, where the easily activated proviruses are unaffected by PADI4, whereas PADI4 and H3cit play a role in chromatin contexts that have an intermediate latency reversal potential.

### Effects of prolonged PADI4 depletion

The K562-derived cell model expresses dCas9-KRAB, making it amendable for CRISPRi. We targeted PADI4 with three gene-specific guide RNAs in addition to a non-targeting control. Interestingly, the main effect of this permanent PADI4 reduction was an increase in spontaneous HIV reactivation (Fig S3). The effect of PADI4-depletion did not result in a reduced HIV-1 activation. However, in PADI4-depleted cells, GSK484 had no effect. Rescuing the PADI4-expression by overexpression of PADI4 reestablished the parental phenotype. These observations suggests that prolonged PADI4 depletion result in compensatory effects by other mechanisms.

### Citrullinated histone H3 at the HIV-1 provirus stimulated latency reversal

We next sought to determine the effect of PADI4-inhibition on chromatin during latency reversal. ChIP using antibodies against H3 and H3cit revealed that T cell activation resulted in an increase of H3cit at the HIV-1 promoter (Fig 3A) and that PADI4 inhibition caused an increase in H3 at the HIV-1 promoter (Fig 3B). In the light of previous research ^17,18^, this was expected. This led us to compare the effect of PADI4 on the HIV-1 promoter using a dedicated chromatin accessibility assay. In addition to more nucleosomes revealed by ChIP, the chromatin at the HIV-1 promoter became more compact after addition of GSK484 alone (Fig 3C). We next compared these results with results from the PLA-ZFP assay where the proviral promoter is marked with an engineered zinc-finger protein (ZFP) that specifically recognizes the HIV-1 LTR ^35,36^. This HIV-1 specific ZFP was introduced into 5A8 cells, to generate the 1C10 cells, with the proviral promoter marked by the protein. As observed using ChIP, the level of H3cit at the HIV-1 promoter was elevated after T cell stimulation (Fig 3D). This effect was less pronounced in the presence of GSK484, confirming a role for *de novo* PADI4-mediated citrullination at the HIV-1 locus during HIV-1 latency reversal. Total histone H3 at the HIV-1 promoter was slightly increased after cell activation (Fig 3E). As expected from previous flow cytometry measurement, GSK484 reduced the GFP signal as observed microscopically (Fig S4A). This technique allows simultaneous detection of H3cit and the effect on HIV-1 activation in single cells. However, due to technical variance, we could not correlate the HIV-1 latency reversal with H3cit levels in individual cells (Fig S4B).

**Fig 3:**
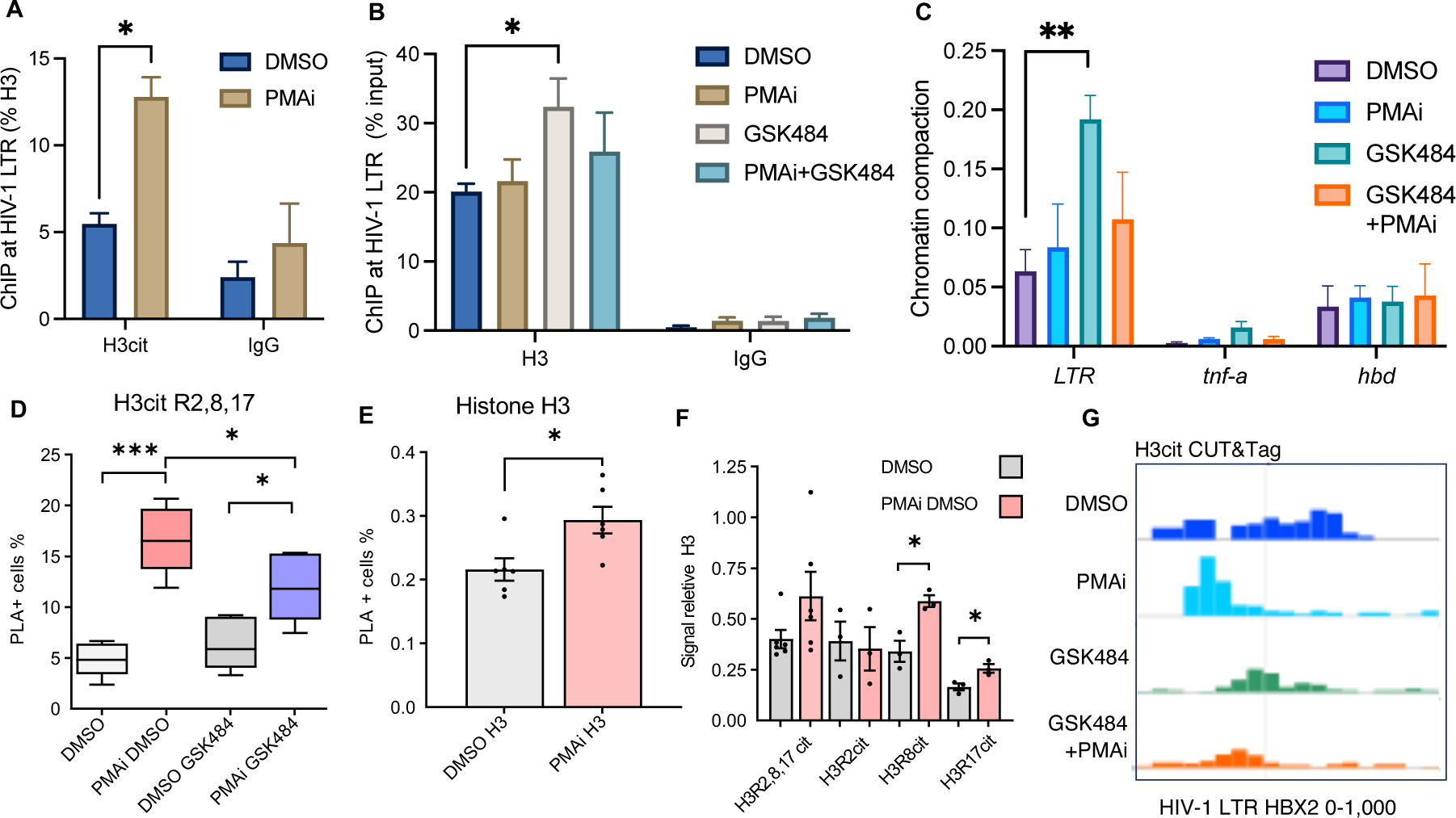
Citrullinated histone H3 at the HIV-1 provirus stimulated latency reversal. (A) H3cit signal at a HIV LTR locus in 5A8 cells, measured by ChIP-qPCR. (B) H3 signal at the HIV LTR locus in 5A8 cells, measured by ChIP-qPCR. (C) Chromatin compaction at the HIV LTR and TNFα. Quantified with chromatin accessibility kit (D) Quantification of H3cit at the HIV-1 LTR in 5A8 cells under PMAi and GSK484 treatment using PLA-ZFP assay (n=6). (E) Quantification of Histone H3 at HIV-1 LTR in 5A8 cells under PMAi treatment using PLA-ZFP assay (n=6). (F) Quantification of different H3cit residues with PLA-ZFP in 5A8 relative to H3 (n=3-6) (G) CUT&Tag seq view of total H3cit after 24 h treatment over the HIV-1 LTR. Grey line indicates TSS. Data is shown as mean ± SEM. Statistical tests shown are unpaired T tests *= <0.05 *= <0.01 *= <0.001

We measured the influence of citrullination at three different H3 arginine residues–H3R2, H3R8 and H3R17. Relative to H3, only H3cit8 and H3cit17 increased significantly, with the strongest relative effect on H3cit8 (Fig 3F).

To map the distribution of H3cit at the provirus in 5A8 cells, we applied CUT&Tag ^37^. Mapping was done using four different conditions, unstimulated or PMAi stimulated cells with or without GSK484 for 24 h. In unstimulated cells, two peaks of H3cit were found at the HIV-1 LTR promoter, coinciding with two well-described nucleosomes (nuc-0 and nuc-1) surrounding the transcription start site (Fig 3G). Upon stimulation, the nuc-1 H3cit signal disappeared. Exposure to GSK484 resulted in weaker signals and less clear nucleosome positioning. PMAi induced activation is known to shift or evict nuc-1 of the HIV-1 promoter ^38^. Inhibition of PADI4-mediated citrullination has previously been associated with nucleosome repositioning ^18^.

### Genome-wide H3cit is stably found at most promoters

The CUT&Tag data provided us with a genome-wide distribution of H3cit in 5A8 cells. Peak calling using MACS2 identified mostly narrow H3cit peaks with a median of 749±20 bp (Fig 4A). Mapping revealed that most (55.4%) H3cit was found at or close to genes (Fig 4B). Aligning the peaks to the transcription start sites of all genes showed that H3cit was predominantly found at promoters, and some in the early coding region (Fig 4C). This global H3cit pattern appears to be stable, as this pattern was not affected by cell stimulation by PMAi or PADI4 inhibition by GSK484 (Fig S5).

**Fig 4:**
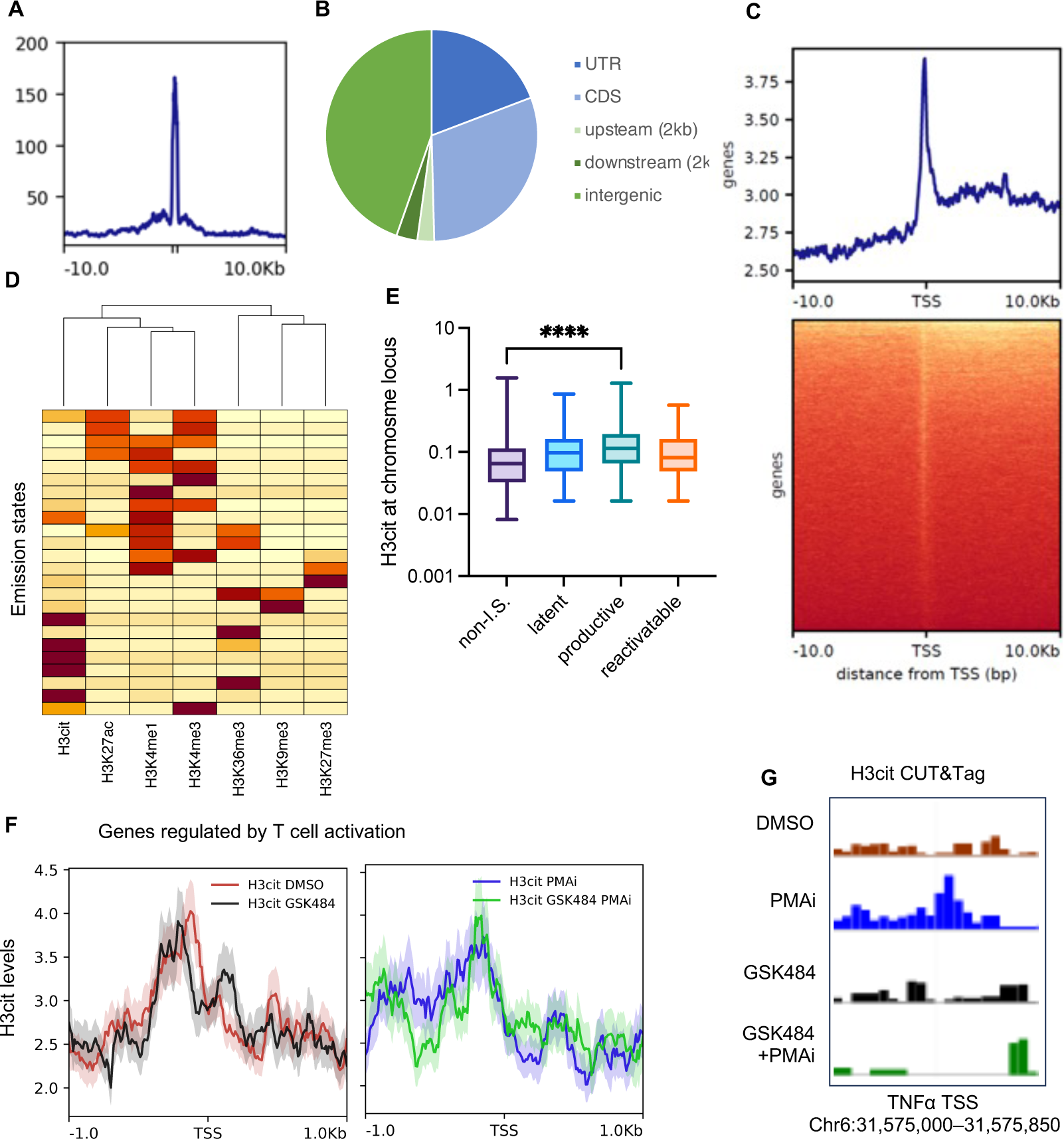
Genome-wide H3cit levels are stable at promoters in 5A8 with HIV-1 preferentially integrating into H3cit chromatin. (A) CUT&Tag peak width profile. (B) Relative enrichment of peaks in genic and non-genic regions. (C) Metagene plot for DMSO treated 5A8 samples (D) ChromHMM analysis showing similarity between chromatin marks. (E) Quantification of H3cit mapped onto known proviral integration sites with corresponding proviral activation. (F) Metagene plot with a gene set regulated by T cell activation (“GSE13738 Resting vs TCR activated CD4 T cell”). (G) Genome browser view of the H3cit signal over the TNFα promoter region. Grey line indicates the TSS.

To relate the H3cit pattern to the distribution of other chromatin marks, we performed a ChromHMM analysis ^39^. As expected, the genome-wide H3cit distribution clustered together with histone marks found at open regions of the genome, such as H3K4me3, H3K27ac and H3K4me1 that are found either at promoters or enhancers. The H3cit pattern matched most closely with that of H3K4me3 (Fig 4D). Both H3cit and H3K4me3 are stable marks, associated with open regions, but not necessarily active gene expression ^40,41^. On the other hand, marks of heterochromatin such as H3K9me3 and H3K27me3 were clustered away from H3cit, as well as H3K36me3, a mark internal to genes where transcription initiation is blocked ^42^.

### Productive HIV-1 integrates in H3cit chromatin

Next, we sought to determine the role of H3cit in initial integration of the HIV-1 sequence. We compared the genome-wide map of H3cit to previous integration site data, stratified on the fate of the provirus ^43^. As expected from H3cit being found in permissive regions of decondensed chromatin, HIV-1 was enriched in regions with H3cit (Fig 4E). Interestingly, H3cit was most often found at integration sites of productive proviruses. Less H3cit was found at latent or latent reactivatable proviruses, implying that the H3cit structure is resistant to silencing, as previously suggested for HERVs ^16^.

### H3cit takes part in activity of T cell receptors genes

No gene set terms were significantly enriched in within the genes associated with H3cit. However, we found 80 genes where new H3cit is formed after PMAi, and 145 genes where the activation-induced citrullination is blocked by GSK484. The overlap was significant as 61 genes were found in both lists (Fisher’s exact test, p<10^-15^). Two clusters of gene sets were significantly enriched here: 1) “T-cell receptor”, “adaptive immunity”, and 2) “citrullination” and “nucleosomes”. Upon closer inspection of these, we noted that for a gene set regulated by T cell activation (“GSE13738 Resting vs TCR activated CD4 T cell” ^44^), GSK484 led to equally distributed H3cit nucleosomes surrounding the TSS. After activation, few changes were observed (Fig 4F). This observation is consistent with the previously observed redistribution of nucleosomes following PADI4-inhibition by GSK484 treatment ^18^. However the PADI4 target seems not to be citrullination of the three H3 residues interrogated here as they remain present.

The TNFα promoter is known to be citrullinated after cell stimulation ^16^. In agreement, we observed *de novo* H3cit at the TNFα TSS after cell stimulation. This effect was completely abolished in the presence of GSK484 (Fig 4G). This demonstrates the possibility of several roles of H3cit in chromatin during T cell activation.

### *De novo* H3cit has a general effect on HIV-1 transcription, but not through Tat or acetylation

To test if the effect on the observed loss of GFP in GSK484 treated cells was a result of reduced transcription or a subsequent process, we extracted RNA from the PMAi induced 5A8 cells. HIV-1 produces both spliced and unspliced transcripts as to create infectious viruses ^13^. We tested the effect of GSK484 on short initiated (5’-LTR), early elongated unspliced (Gag), multiply spliced (Tat-Rev), and late elongated (GFP) transcripts. These transcript types were equally reduced by 30% after GSK484 treatment, although significance was only reached for GFP and Tat-Rev (Fig 5A).

**Fig 5:**
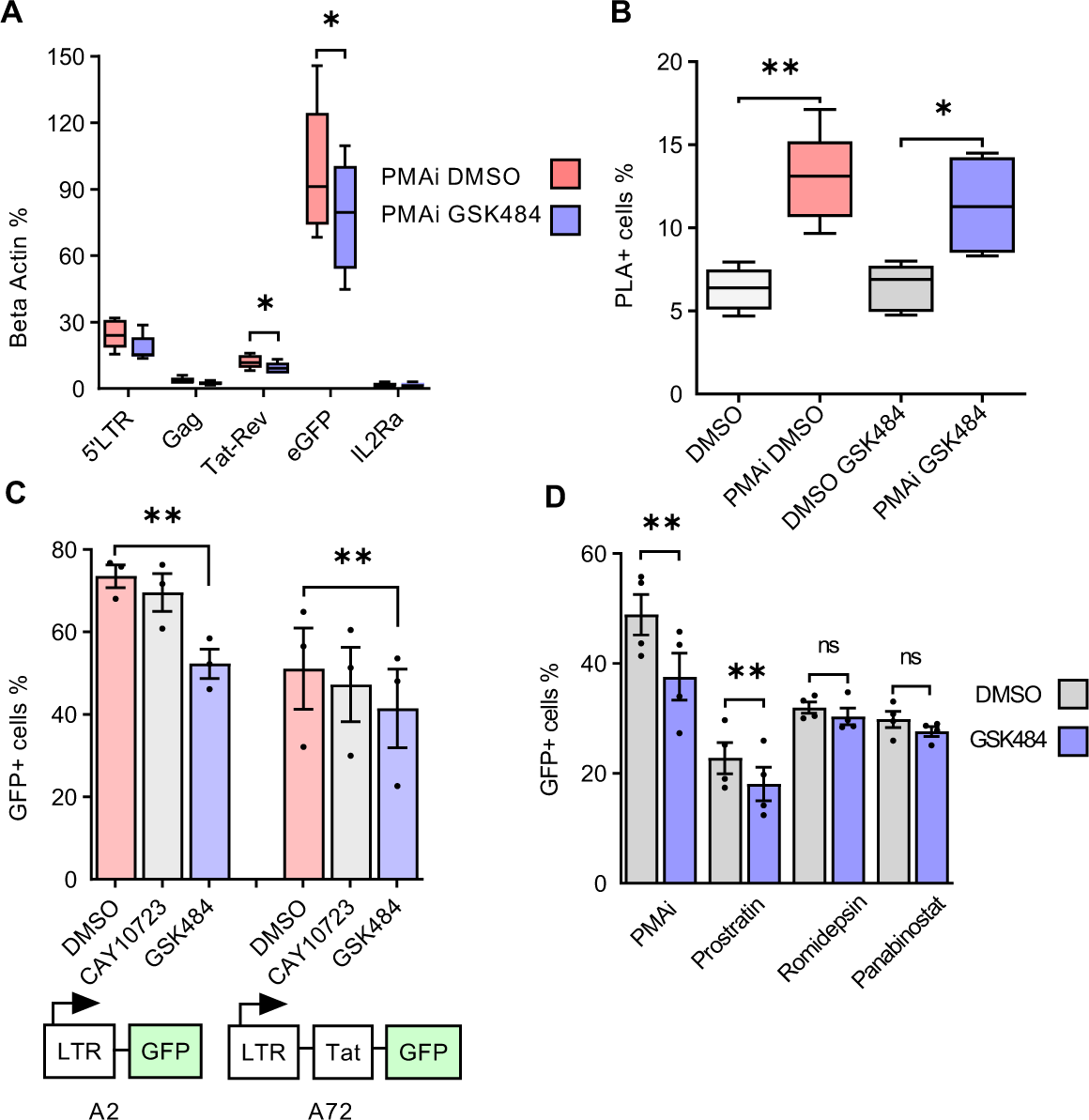
*de novo* H3cit has a general effect on HIV-1 transcription, but not through Tat or acetylation. (A) RT-qPCR quantification of short initiated (5’-LTR), early elongated unspliced (Gag), multiply spliced (Tat-Rev), and late elongated (GFP) transcripts in 5A8 treated with PMAi and GSK484. Statistical tests shown are paired T tests *p<0.05 (n=5). (B) Levels of Tat at the HIV-1 promoter in 1C10 cells with PMAi and GSK484 using PLA-ZFP assay. Statistical tests shown are unpaired T tests * p<0.05, ** p<0.01, *** p<0.001 (n=6). (C) HIV-1 transcription of in Tat negative J-Lat A2 and Tat positive A72 cells treated with PMAi and PADI inhibitors, measured by flow cytometry. Statistical tests shown are paired T tests * p<0.05, ** p<0.01, *** p<0.001 (n=3). (D) 5A8 cells treated with PMAi and other latency reversal agents and GSK484 (n=4). Data is shown as mean ± SEM.

We next tested if H3cit impaired Tat function. We used PLA-ZFP to detect recruitment of Tat to the LTR in 1C10 cells. Tat was still recruited to the LTR after GSK484 exposure (Fig 5B). Normally Tat is responsible for augmenting the transcription. However, in single cells, the amplitude of GFP expression was not affected by GSK484, only the amount of GFP+ cells, further suggesting no link between Tat and PADI4. To confirm the independence of Tat, we compared J-lat cells with partial HIV-1 genomes, either LTR-GFP (A72), or LTR-Tat-GFP (A2) ^31,45^. Both cell lines were exposed to PADI-inhibitors during PMAi stimulation and showed similar levels of GFP reduction after both Cl-amidine and GSK484 (Fig 5C). This is consistent with a Tat-independent mechanism.

To further explore which host mechanisms are involved in PADI4-mediated HIV-1 transcription, we tested the effect of GSK484 with different latency reversal agents. In addition to PMAi, latency reversal by the protein kinase C agonist Prostratin was affected by GSK484. Latency reversal by HDAC inhibitors Romidepsin and Panabinostat was not affected by GSK484 (Fig 5D). This suggests that the observed effect of PADI4 is either independent from histone acetylation, or that the subset of cells in which PADI4 plays a role for HIV-1 transcription does not overlap with the one activated by HDACi.

### Heterochromatin forms over the HIV-1 provirus in the absence of H3cit

Given our finding that *de novo* citrullination of H3 and possibly other targets appears to prevent HIV-1 transcription in primary cells from PLWH as well as in several model systems, together with the previous finding of H3cit preventing heterochromatin over HERVs ^16^, we set out to study the impact of H3cit on heterochromatin formation over the HIV-1 provirus. The 5A8 cells have the provirus integrated in a permissive region, and the proviral chromatin is predominantly associated with active histone marks ^35^. This chromatin structure mirrors the majority of proviral integration events, as HIV-1 tends to target open chromatin ^46^. However, once ART is initiated, new infections are prevented. The immune system targets cells producing viral proteins, resulting in an advantage of latently infected cells. Long-term ART results in the provirus being found in epigenetically silent regions ^6^. To replicate this epigenetic state, we modified the 5A8 cells to silence the provirus. We tethered a KRAB (Krüppel Associated Box) domain to the engineered ZFP that specifically recognizes the HIV-1 provirus used for the PLA experiments ^35^. We introduced this KRAB-ZFP protein into the cells and selected for monoclonal cell lines. We characterized two clones in particular, 2C10 with the ZFP-KRAB, and 1C10–the control cell line with only the ZFP construct. In the 2C10 cells, H3K9me3 at the promoter was confirmed by ChIP (Fig 6A). The HIV-1 inducibility after 24 h PMAi stimulation was lower in 2C10 compared to the control 5A8 clone 1C10 (Fig 6B). Among the cells that still had inducible HIV-1, GSK484 further reduced the reactivation. As expected from less HIV-1 reactivation, the recruitment of Tat was less significant in the 2C10 cells (Fig 6C). This shows a limited effect of H3cit in established heterochromatin.

**Fig 6:**
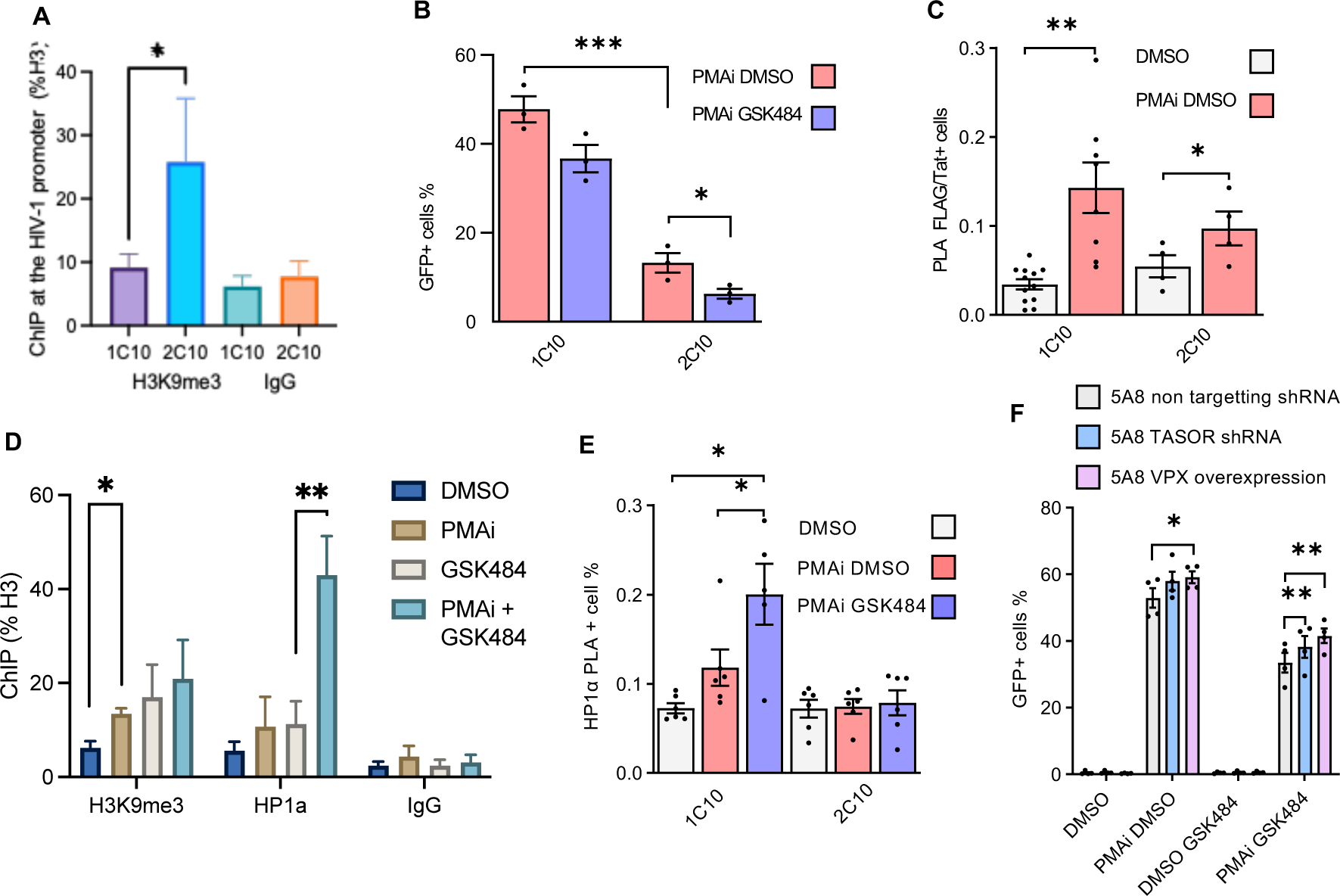
Heterochromatin forms over the HIV-1 provirus in the absence of H3cit. (A) H3K9me3 at the HIV-1 locus in 1C10 (ZFP) and 2C10 (ZFP-KRAB) cells measured by ChIP-qPCR. Statistical tests shown are unpaired T tests * p<0.05 (n=3). (B) 1C10 and 2C10 cell HIV-1 transcription with PMAi and GSK484 treatment. Statistical tests shown are unpaired T tests * p<0.05, ** p<0.01, *** p<0.001 (n=3). (C) Tat proximity to HIV-1 promoter in 1C10 and 2C10 cells with PMAi treatment. Statistical tests shown are paired T tests * p<0.05, ** p<0.01 (n=6-12). (D) H3K9me3 and HP1α levels at the readth locus in 1C10 measured by ChIP-qPCR (E) HP1α proximity to the HIV-1 promoter in 1C10 and 2C10 cells with PMAI and PMAi and GSK484 treatment (n=5-6). (F) GFP quantification by flow cytometry of 5A8 TASOR RNAi knockdown and 5A8 VPX overexpression cells (n=3). Data is shown as mean ± SEM.

Next we wanted to investigate if H3cit prevents heterochromatin development. We hypothesized that *de novo* citrullination of the HIV-1 promoter prevents HP1α binding and functional heterochromatin formation, despite occurrence of H3K9me3. From a previous study ^35^, we know that following activation H3K9me3 levels increase at the HIV-1 provirus. We confirmed this finding using ChIP (Fig 6D). However, H3K9me3 is not sufficient to establish heterochromatin. To form functional heterochromatin, the H3K9me3 should be enclosed in HP1α. In HERVs, H3cit8 hinders HP1α binding to H3K9me3 ^16^. Consequently, by activating the cells and preventing citrullination by GSK484, HP1α would be given a platform to bind. We tested the recruitment of HP1α to the HIV-1 promoter by both ChIP (Fig 6D) and PLA (Fig 6E). Consistently using both techniques, HP1α binds specifically to HIV-1 after cell stimulation when PADI4 is inhibited. In the 2C10 cells, where the provirus was embedded in heterochromatin before the T cell stimulation, HP1α was not found above background levels (Fig 6E).

### H3cit partly inhibited HUSH-mediated chromatin silencing

New heterochromatin formation over foreign DNA can be directed by the HUSH complex ^47^. To test if our observations can be explained by HUSH, we reduced expression of HUSH component TASOR by shRNA, as well as ectopically expressed VPX that is known to repress HUSH ^48^. In 5A8 cells, the latency reversal effect of GSK484 after T cell activation depended significant on HUSH (Fig6F). However, only part of the effect (approx. 20%) could be explained by HUSH.

## Discussion

In this study, we demonstrated that *de novo* PADI4-mediated citrullination of histone H3 at the HIV-1 promoter inhibits the formation of heterochromatin and promotes HIV-1 transcription following cell activation. When PADI4 activity was blocked during cell activation, new citrullination of proviral nucleosomes did not occur, allowing stable heterochromatin to form. The HUSH complex was partly involved in the heterochromatin formation, but the main effect appeared to come from an unidentified source. Proviruses within citrullinated chromatin were resistant to epigenetic silencing. This mechanism is particularly relevant for HIV-1 proviruses integrated in permissive chromatin regions, which are less prone to latency and more responsive to latency reversal agents. Inhibiting *de novo* H3cit induces chromatin compaction at the HIV-1 promoter. We also provide the first genome-wide map of H3 citrullination in T cells, showing that this mark is enriched at promoters and associated with permissive chromatin.

### Implications for the reservoir in people living with HIV-1

We have uncovered a mechanism that explains one aspect of the differences in the HIV-1 reservoir between chronic viremia and after long term ART in PWH. Cells continuously producing viruses undergo cytopathic effects and are targeted by the immune system ^7,49^. We observe a mechanism that is active in viremic individuals but contributes less after long term ART. The latent reservoir is, at least in part, transcriptionally active ^1,3,28,50^. Our study highlights the heterogeneity of the reservoir, particularly during chronic infection before ART. When ART is initiated, pressure to induce reactivatable latent proviruses increases. The intact viruses date to the time of ART initiation ^51^. The state of the reservoir when ART is initiated is key for post-treatment control and a potential HIV-1 cure^52^. Over time an increasing fraction of the intact provirus is found in heterochromatin ^6^. Part of the reservoir is susceptible to histone acetylation and can be cleared even under ART ^53^. Future studies are required to reveal the fate of the cells with citrullination at the promoter, that cannot be epigenetically silenced after ART initiation. Many of these cells are likely eliminated through cytopathic effects or immune detection, unless they are cytotoxic CD4 T cells which resist cell death^50^. A subset of proviruses might become latent solely as a consequence of the cell entering a resting state ^54^, and then stochastic events cause reactivation ^55^. This points to the possibility of reducing the fraction of the reservoir that is subject to stochastic reactivation by manipulating PADI4 during initiation of ART.

### Epigenetic interplay of H3cit and other PADI4 targets

H3cit is linked to active transcription and open chromatin across various cellular contexts ^16,25–27^. In addition to H3, PADI4 targets numerous proteins in the nucleus, including histone H1 and HP1γ ^25,27^. We cannot exclude that modifications of these proteins also play a role in the phenotype we observe. Here we detailed the role of H3cit8. Notably, other H3 residues can also be citrullinated, potentially influencing the cell differently. The H3cit26 mark is recognized by SMARCD1, that has been suggested as a link between H3cit and heterochromatin ^26^. In mice ES cells, H3cit26 and H3K9me3 do not colocalize at basal levels. However, after PADI inhibition, H3K9me3 levels appear at the sites of H3cit26^26^. Arginines, which can be citrullinated, are often found adjacent to lysines, key residues for epigenetic modifications. Moreover, arginine methylation and citrullination are mutually exclusive ^20^. Both arginine methylation and citrullination are stable histone modifications and whereas a short pulse of PADI4 inhibition is unlikely to shift the balance between methylated and citrullinated arginine residues, prolonged reduced levels of PADI4 could have this effect ^18^. Arginine methylation of H3 through CARM1 has a known repressive effect on HIV-1 latency ^56^. We conclude that PADI4 plays a balancing role in HIV-1 transcription, depending on both the cell type and the site of proviral integration.

### Limitations of the study

Our study is confined to CD4 T cells from peripheral blood, despite the presence of the HIV-1 reservoir at several other anatomical sites in the body. While our findings are grounded in clinical samples, the mechanistic insights primarily derive from cell models. We were unable to track PADI4-depleted cells as they reverted to latency to determine if they were prevented from subsequent reactivation. To enhance the generalizability and applicability of the findings, we employed several different cell models and complementary techniques. However, we were unable to test our hypothesis *in vivo* or in animal models. The small sample size in parts of the test is a limitation for the generalizability.

### Conclusions

Our findings reveal a novel epigenetic mechanism that regulates HIV-1 transcription and latency, suggesting that PADI4 could serve as a potential therapeutic target for interventions aimed at reducing the size of the latent reservoir required for a future HIV-1 cure.

## Methods

### Cell Culture and isolation

J-lat 5A8, 1C10, K562 and all primary cells were cultured in cytokine-free media (RPMI 1640 medium (Hyclone, Cat# SH30096_01), 10% FBS (Life Technologies, Cat# 10270–106), 1% Glutamax (Life Technologies, Cat# 35050), 1% Penicillin-streptomycin (Life Technologies, Cat# 15140–122)).

Primary cells were obtained from 50 ml of fresh blood from study participants (Ethical permits 2017/1138-31/4 and 2018/102-31/1). Blood samples were diluted with an equal amount of PBS and carefully overlayed on Ficoll-Paque Plus solution (Cytiva, Cat# 17144003). Centrifugation was conducted at RT and 400 g for 30 minutes without brake. Resting CD4 T cells were isolated from the obtained PBMCs by a two-step magnetic isolation using the Miltenyi Biotec magnetic associated cell sorting (MACS) platform. For the first step, the CD4 negative cell isolation kit (Cat# 130-096-533) was used according to the manufacturer’s protocol. Secondly, resting CD4 T cells were negatively selected with magnetic beads against the activation markers CD25, CD69 and HLA-DR.

### Cell activation

CD4 T cells isolated from peripheral blood and immediately put in *ex vivo* culture. After 16 h resting, cells were exposed to PADI4 inhibitor GSK484 for 3 h followed by T cell activation by phorbol-12-myristate-13-acetate and ionomycin (PMAi) for 21 h. Both DNA and RNA were extracted from the cells, and culture supernatant RNA was extracted to quantitate release of viral particles. DNA was extracted either by Allprep (Qiagen, Cat#80204) or QIAamp DNA Mini Kit (Qiagen, Cat#51304). Viral RNA was isolated by QIAamp Viral RNA Mini Kit (Qiagen, Cat#52904). The DNA was used to measure the HIV-1 reservoir using IPDA ^29^. For cell associated RNA, the LTR region and the multiply spliced RNA was investigated separately. Cell-free RNA was measured by the amount of intact virus RNA in the supernatant ^30^.

### Chemicals to induce proviral activation

Cells were exposed to latency reversal agents as indicated with the following concentrations used: Phorbol 12-myristate 13-acetate (PMA, Sigma-Aldrich Cat# 79346) final concentration 50 ng/ml, ionomycin (Sigma-Aldrich Cat# I0634) final concentration 1 µM. Ingenol-3-angelate PEP005 (Sigma-Aldrich, Cat# SML1318) final concentration 12 nM, panobinostat (Cayman Chemicals, Cat# CAYM13280) final concentration or 150 nM, JQ1 (Cayman Chemicals, Cat#CAYM11187) final concentration 100 nM, prostratin (Sigma-Aldrich, Cat#P0077) final concentration 6 μM.

PADI4 inhibitor GSK484 (MCE HY-100514) was added, if applicable, at a final concentration of 10 µM. Pan PADI inhibitor Cl-Amidine (Cayman Chemicals. Cat#CAYM10599) was added at a final concentration of 200 µM. Inactive GSK484 and GSK199 control GSK106 (MCE HY-120343) was added at a final concentration of 10 µM. Padi4 Inhibitor GSK199 (MCE HY-103058) was added at a final concentration of 10 µM

### Flow cytometry

Cells were stained with LIVE/DEAD Fixable Violet Dead Cell Stain (Thermo Scientific, Cat# L34955) and fixed in 2% formaldehyde for 15 min. if required Flow analysis was performed on a CytoFLEX S (Beckman Coulter). Individual flow droplets were gated for lymphocytes, viability, and singlets. Data were analyzed by Flowjo 10.1 (Tree Star).

When staining for CD25 and CD69. Cells were washed with PBS (Gibco #18912014) + 0.5% BSA (FACS buffer) then incubated with 1/100 CD25 (BD, Cat#555434) and CD69 1/20 (BD, Cat# 560711) antibody diluted in FACS Buffer for 30 min at 4°C. Cells were then washed with 100ul of FACS buffer then resuspended in 100ul FACS buffer and 1/1000 dilution of Live/dead stain (Thermo Scientific, Cat# L34964) and incubated for 15 min at 4°C. Cells were washed in 100 µl FACS buffer then resuspended in 2% formaldehyde PBS for 15 min in the dark at room temperature. Finally, cells were resuspended in 100 µl of ice-cold PBS prior to flow cytometry.

When staining for KI-67 cells were washed twice with PBS and resuspended by vortexing while adding ice cold 70% Ethanol. This was followed by a 1 h incubation at −20°C, three washed with FACS buffer then resuspension in FACS buffer at 1−10^6^ cells/ml. 5 µl of APC anti-human Ki-67 Antibody (BioLegend, CAT# 350513) in 100 µL of cell solution. This was incubated for 30 min in the dark at room temperature, washed with FACS buffer then resuspended in 100 µl of ice-cold PBS prior to flow cytometry.

### KRAB domain expression in 5A8 cells

The KRAB domain from pHR-SFFV-dCas9-BFP-KRAB (Addgene plasmid #46911) was cloned into plasmid pHR-SFFV-ZF3-P2A-BFP using Gibson assembly (New England Biolabs Cat#E2611S) by substituting the KRAB region in between the ZFP3 and P2A region.

The contruct was co-transfected with psPAX and pMD2.G into HEK293T cells using lipofectamine LTX (Invitrogen Cat# 15338030) according to the manufacturer’s instructions. 20 µL of the viral supernatant was added to 180 µL of K562-Lat cell culture in a round bottom 96 well plate and spinoculated for 300g for 1hr at room temperature. These were puromycin selected with 1.5 µg/ml puromycin RPMI media for 5 days.

### Immunoblotting

Cell pellets were lysed in lysis buffer (150 mM NaCl, 50 mM Tris, 1% Triton-X100, 1 mM orthovanadate). Protein concentrations were calculated using Bradford assay (Bio-Rad Cat#5000006). Human Histone H3 (citrulline R2 + R8 + R17) peptide (Abcam, Cat# ab32876) was used in a competition assay. Samples were diluted to 1× Laemmli buffer (Bio-Rad, Cat# 1610747) with 10 mM DTT. Samples were heated for 10 min at 70°C and then loaded on a 12% Mini-protean TGX gel (Bio-Rad Cat# 4561044). The gels were run for approximately 1 h at 100 V. Proteins were transferred onto a PVDF membrane (Bio-Rad, Cat# 1704156) and blocked in 5% skimmed milk in PBS-T for 1 h at room temperature. Afterwards, the membrane was incubated at 4°C overnight with primary antibodies (1:1000) in 1% milk in PBS-T. We used primary antibodies as above. After washing and addition of secondary HRP-conjugated antibodies, membranes were developed using Pierce ECL Plus Western Blotting Substrate (Thermo Scientific Cat# 32132). Membranes were stripped (Restore plus Western blot stripping buffer Thermo Scientific Cat# 46430) and re-probed using a second set of control primary antibodies. Blots were run and visualized using iBright FL1500 Imaging System (ThermoFisher).

### Chromatin immunoprecipitation

ChIP-qPCR was performed using the iDeal ChIP-qPCR protocol (Diagenode, Cat# C01010180). Each ChIP reaction was performed on 2 × 10^6^ cells. Cells were fixed with 11% formaldehyde for 10 min in room temperature. Sonication was performed at 30 s in eight cycles (Bioruptor Pico, Diagenode, Cat# B01060010). ChIP was performed using antibody. ChIP eluates were purified with Wizard SV Gel and PCR clean-up system (Promega, Cat# A9282). Primer sequences are shown in Table S2. PCR reactions were performed with Powerup SYBR green master mix (2×) (ThermoFisher, Cat#A25742) using 40 cycles on an Applied Biosystems 7500 Fast Real-Time PCR System (ThermoFisher).

### CUT&Tag seq

Version 3 CUT&Tag protocol (Kaya-Okur and Henikoff, 2020) was used as described. A 1/400 dilution of Anti-Histone H3 (citrulline R2 + R8 + R17) antibody (Abcam, Cat# ab5103) was used per sample. Thirteen cycles of PCR were used for amplifying the libraries. Reads were trimmed, then, mapped onto the GB38 human reference genome with Bowtie2.

### Gene set enrichment

Genes with log2 ratio>0.5 difference between DMSO and PMAi, or PMAi and PMAi, GSK484 were selected (Table S3). The lists were analyzed using the DAVID tool ^57^.

### Proximity ligation assay (PLA)

Cells were adhered to coated polylysine coated microscope slides (VWR, Cat#6310107) marked with a hydrophobic barrier using A-PAP pen (Histolab, Cat# 08046N) and a modified PLA protocol was followed. In short, cells were washed with PBS and allowed to settle onto the polylysine slides). PLA was performed according to the manufacturer’s protocol (Sigma-Aldrich, cat#Duo92007) with a few modifications: PLA plus and minus probes were diluted 1:20, amplification buffer (5×) was used at 10×. All washes were performed in PBS. Antibodies (1:1,000) were used against Tat (Abcam Cat#43014), FLAG M2 (Sigma-Aldrich Cat# F1804 lot SLCF4933), H3 (Abcam, Cat# ab1791), H3R2cit (Abcam, Cat# ab176843), H3R8cit (Abcam, Cat# ab219406), H3R17cit (Abcam, Cat# ab219407), H3R2cit+R8cit+R17cit (Abcam Cat# ab5103). Before DAPI staining (Life technologies, Cat# 62248) and mounted with mounting medium with DAPI (Sigma-Aldrich, Cat# DUO82040), FITC-conjugated anti-GFP (Abcam, Cat#ab6662) was applied (1:1000) for 1 hour at ambient temperature protected from light. Coverslips (VWR, Cat# MARI0117530) were sealed onto slide with nail polish and stored at 4 °C overnight before imaging. Slides were imaged using a Panoramic Midi II slide scanner (3DHistech) and images were exported using the CaseViewer application. Images were analyzed with ImageJ (version 2.0.0-rc-69/1.52) and macros developed in house. Each RGB image was separated into separate channels. Based on the blue (DAPI) channel, nuclei were identified. In the red channel, spots (“maxima”) were detected with thresholds 6–48.

### Quantitative PCR (qPCR)

RNA was extracted from cells using RNeasy plus kit (Qiagen Cat#74134) according to the manufacturers instructions. Then cDNA was made using Revertaid as described (Thermo Scientific Cat#K1621). The cDNA was diluted 1/10 in distilled water and used with 400 nM of each primer (final concentration) and SYBR Green Master Mix for qPCR (Fisher Scientific Cat# A25742) in a Microamp Fast Optical 96 Well Reaction Plate (Fisher Scientific Cat#4346906). The PCRs were run using Applied Biosystems 7500 Fast Real-Time PCR System with 40 cycles.

### Digital droplet PCR (ddPCR)

ddPCR was performed with the QX200 Droplet Digital qPCR System (Bio-Rad). Each reaction consisted of 20 μL, containing 10 μL Supermix for Probes without dUTP (Bio-Rad, Cat# 1863024), 900 nM primers, 250 nM probe (labelled with HEX or FAM), and 8 μL undiluted cellular DNA. RT-ddPCR was used according to ^30^, using 11 μL undiluted RNA and the One-Step RT-ddPCR Advanced Kit for Probes (Bio-Rad, Cat# 1864021). Droplets were generated. Emulsified PCR reactions were performed with a C1000 Touch thermal cycler (Bio-Rad), with the following protocol: 95 °C for 10 min, followed by 40 cycles of 94 °C for 30 s and 57 °C for 60 s, and a final droplet cure step of 10 min at 98 °C. Each well was then read with a QX200 Droplet Reader (Bio-Rad). Droplets were analysed with QuantaSoft version 1.5 (Bio-Rad) software in the absolute quantification mode.

### Data availability

The datasets generated in this study is publicly available through the Gene Expression Omnibus (GEO) GSE273899

## Acknowledgements

We are most grateful to the study participants without whom this study would not have been possible. We would also like to thank BEA, MedH flow facility, as well as Mukesh Varshney and Zhichang Huang for technical assistance. Funding: This study was supported by grants from CIMED FoUI-973749, VR 2019–00991, KI 2019-00846, Cancerfonden 190412Pj (to J.P.S.), Swedish Physicians Against AIDS Research Foundation FOb2020-0004 (to J.P.S), FOa2022-0001, FOa2023-0003 (to B.B.J.), KI Foundation for Virus Research 2024-00634 (to L.L.).

## Author contributions

Conceptualization, L.L., J.P.S.; investigation and validation, L.L, B.B.J., B.L., H.R., J.P.S.; clinical recruitment, P.N., O.K.; supervision, J.P.S.; funding acquisition, J.P.S. and B.B.J. Writing – original draft, L.L., B.B.J., and J.P.S.; review & editing, all co-authors.

