## Supplementary figures for "PADI4-mediated citrullination of histone H3 stimulates HIV-1 transcription"

### Supplementary material Love et al

**Table S1:** Study participant characteristics

**Table S2:** Genes with log2 ratio>0.5 difference between DMSO and PMAi, or PMAi and PMAi + GSK484

**Table S3:** Primer sequences for ddPCR and ChIP.

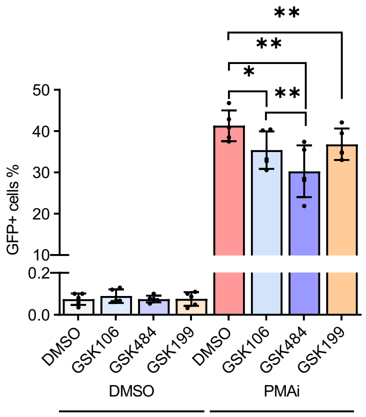


**Fig S1:**

5A8 treated with PADI4 inhibitors for 24 h with GSK484, GSK199 and inactive control GSK106, GFP measured by flow cytometry (n=4-5).

(B)

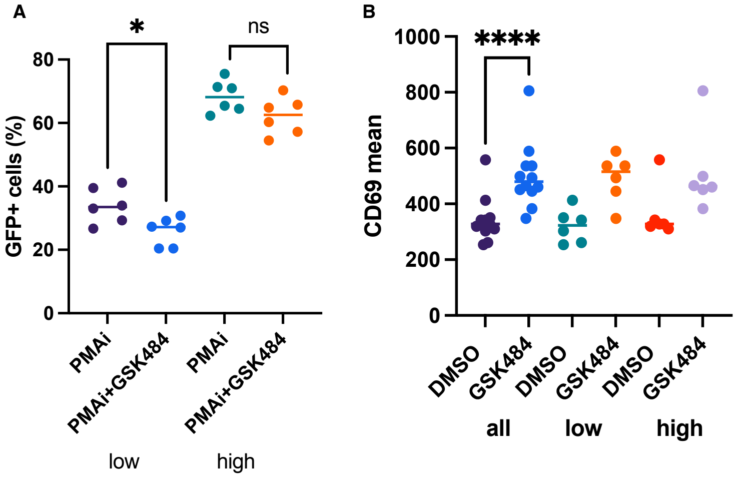


**Fig S2:**

(A) GFP quantification of monoclonal selected heterozygous PADI4 5A8 cells treated with PMAi and GSK484. Stratified into high and low GFP groups. GFP measured by flow cytometry, (n=6)

(B) CD69 quantification of monoclonal selected heterozygous PADI4 5A8 cells in A. Stratified into high and low GFP groups. CD69 measured by flow cytometry. (n=6)

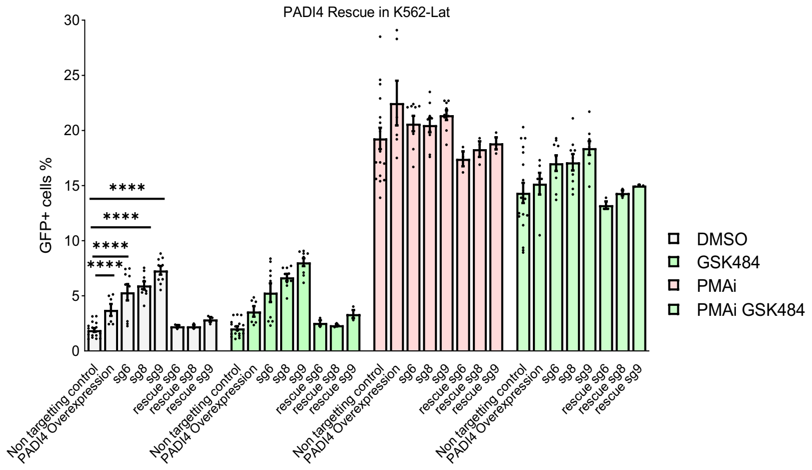


**Fig S3:**

GFP quantification of PADI4 knockdown and rescue in K562-Lat cells treated with PMAi and GSK484. (n=3-14)

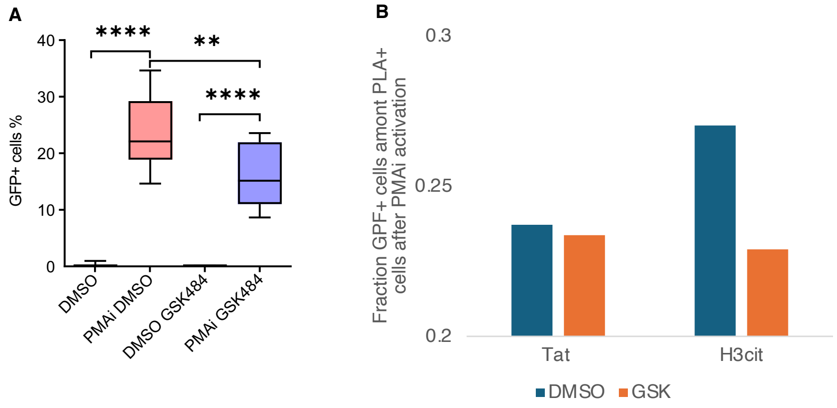


**Fig S4:**

(A) 5A8 treated with PMAi and GSK484. GFP measured by fluorescent microscopy as part of PLA (n=11).

(B) Fraction of GFP+ cells with PLA-Tat signal from PMAi treated 1C10 cells (n=5).

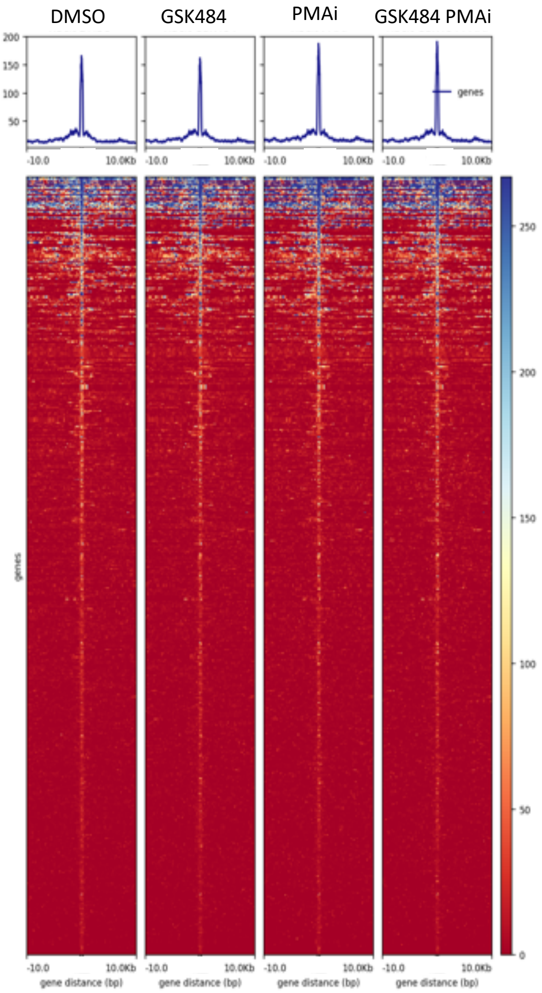


**Fig S5:**

Metagene plot for H3cit peaks in DMSO, GSK484, PMAi and PMAi/GSK484 treated conditions in 5A8.
